# Water inside the selectivity filter of a K^+^ ion channel: structural heterogeneity, picosecond dynamics, and hydrogen-bonding

**DOI:** 10.1101/2023.11.16.567415

**Authors:** Matthew J. Ryan, Lujia Gao, Francis I. Valiyaveetil, Alexei A. Kananenka, Martin T. Zanni

**Affiliations:** Department of Chemistry, University of Wisconsin-Madison, Madison, WI 53706, USA; Department of Chemical Physiology and Biochemistry, Oregon Health & Science University, Portland, OR 97239, USA; Department of Physics and Astronomy, University of Delaware, Newark, DE 19716, USA

## Abstract

Water inside biological ion channels regulates the key properties of these proteins such as selectivity, ion conductance, and gating. In this Article we measure the picosecond spectral diffusion of amide I vibrations of an isotope labeled KcsA potassium channel using two-dimensional infrared (2D IR) spectroscopy. By combining waiting time (100 - 2000 fs) 2D IR measurements of the KcsA channel including ^13^C^18^O isotope labeled Val76 and Gly77 residues with molecular dynamics simulations, we elucidated the site-specific dynamics of water and K^+^ ions inside the selectivity filter of KcsA. We observe inhomogeneous 2D lineshapes with extremely slow spectral diffusion. Our simulations quantitatively reproduce the experiments and show that water is the only component with any appreciable dynamics, whereas K^+^ ions and the protein are essentially static on a picosecond timescale. By analyzing simulated and experimental vibrational frequencies, we find that water in the selectivity filter can be oriented to form hydrogen bonds with adjacent, or non-adjacent carbonyl groups with the reorientation timescales being three times slower and comparable to that of water molecules in liquid, respectively. Water molecules can reside in the cavity sufficiently far from carbonyls and behave essentially like “free” gas-phase-like water with fast reorientation times. Remarkably, no interconversion between these configurations were observed on a picosecond timescale. These dynamics are in stark contrast with liquid water that remains highly dynamic even in the presence of ions at high concentrations.

## INTRODUCTION

Water exhibits diverse roles in molecular cell biology.^1, 2^ Water in the vicinity of a protein surface is called “hydration water”. The properties of protein hydration water deviate markedly from those of the bulk^3-7^ and can significantly modulate the structure and function of proteins.^1, 2, 4, 8-11^ That makes the properties of water near or within biomolecules, and the extent to which they deviate from the bulk, a topic of much interest.^3-7^ In bulk liquid water the reorientation relaxation is fast, with an exponential relaxation timescale of 2.6 ps, which occurs through concerted hydrogen bond rearrangement.^3, 5, 12^ In the proximity of biomolecules, where the hydrogen-bond network is disrupted, the reorientation relaxation of water becomes 2-5 times slower and is nonexponential due to a broad distribution of reorientation times.^4, 13-19,19^ Water molecules in deep crevices and pockets in the interior of a protein exhibit even slower dynamics reaching nanosecond timescales and longer.^4, 7, 8, 18, 20-22^ Strongly bound localized water molecules reorient on millisecond timescales. Such immobile water can be detected using high-resolution X-ray crystallography and cryo-electron microscopy at low temperature.^8, 23, 24^ These crystallographic water molecules may have functional roles.^7^ In contrast, fluctuating water molecules cannot be detected with certainty using X-ray crystallography.

Cation-selective ion channels are an example of proteins whose properties such as conductance, selectivity, and the physiological state are controlled by water.^25-34^ The KcsA potassium ion (K^+^) channel from *Streptomyces lividans* has been integral in providing the structural basis for understanding the function of K^+^ channels.^27, 35^ It has high sequence homology and functional similarity to mammalian potassium channels.^36^ The high-resolution structures of the KcsA channel reveal ion binding sites in the extracellular side, in the selectivity filter, and in a water-filled cavity adjacent to the selectivity filter (Figure 1).^26,35, 37^ In crystal structures of KcsA, crystallographic water molecules are found behind the selectivity filter, on the extracellular side and in the central cavity.^38, 39^ Water is present in the selectivity filter as well and it participates in K^+^ permeation. Therefore, it is fundamentally important to understand the behavior of water in the selectivity filter.

**Figure 1.**
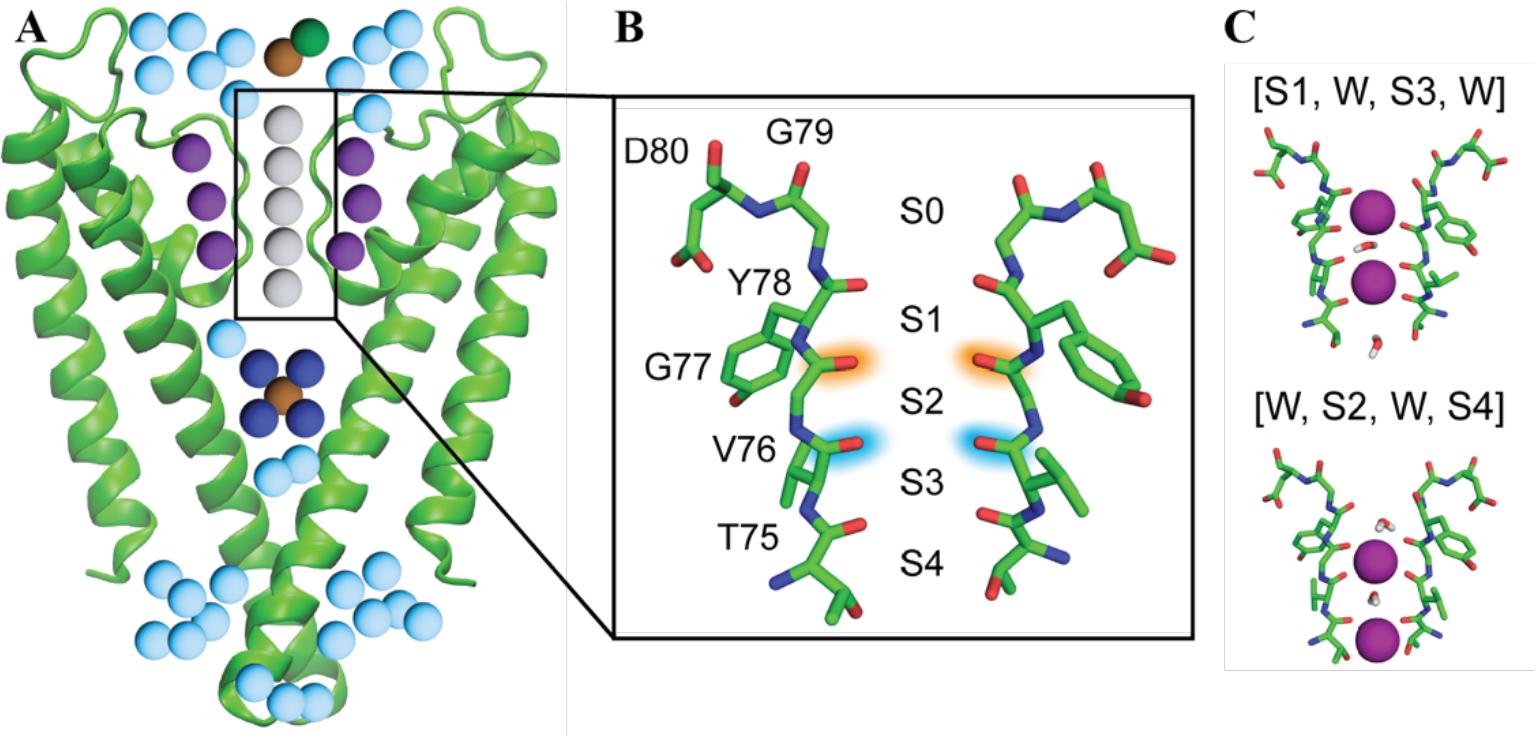
(A) Schematic illustration of the pore domain of the KcsA channel with closed intracellular gate and a conductive selectivity filter. Note that only two of the four protein chains are shown for clarity. Shown are the K^+^ ions (brown), crystallographic water coordinating K^+^ ion in the central cavity (dark blue spheres) and on the extracellular side (dark green sphere), behind the selectivity filter (purple spheres), mobile water (light blue spheres), and the five binding sites inside the selectivity filter (gray spheres) each accommodating either a water molecule or K^+^ ion. (B) The selectivity filter. Isotope-labeled residues Val76 and Gly77 whose 2D IR spectra measured and analyzed in this work are highlighted in blue and orange, respectively. (C) The two configurations in the selectivity filter considered in this work. K^+^ ions shown in purple.

The selectivity filter, being only ∼4.5 Å wide, is the narrowest part of the channel and is responsible for the high selectivity of K^+^ channels. It is formed by segments of the four subunits with the highly conserved sequence Thr75-Val76-Gly77-Tyr78-Gly79 (TVGYG) (Figure 1B).^27, 38^ Five binding sites, S0-S4, are formed by the backbone carbonyl groups from this sequence together with the threonine hydroxyl group.^27, 38, 40-45^ K^+^ ions are tightly coordinated in the selectivity filter: ion binding sites are 3.5 Å apart and the average distance between a K^+^ ion and a carbonyl oxygen atom is 2.85 Å.^42^ The extreme narrowness of the selectivity filter also creates a unique confinement for a single water molecule whose oxygen atom is closer to the protein than the thickness of water’s first coordination shell (*ca*. 3.5 Å) which is also approximately the spatial extent of the first hydration layer.^4^

Very little is known about the dynamics of water in ion channels.^25, 46, 47^ Nuclear magnetic resonance spectroscopy (NMR) can be used to study protein and water dynamics on nanosecond to millisecond timescales.^29, 48-51^ However, because water in the selectivity filter is mobile, one might expect that its dynamics occurs on faster timescales. In this Article we use two-dimensional infrared (2D IR) spectroscopy to study picosecond dynamics of water in the selectivity filter of the KcsA channel.^52, 53^ 2D IR is an intrinsically structure-specific method that measures molecular vibrations. We focus on the amide I vibration which is primarily a carbonyl stretching motion of the peptide group.^54-58^ Because the amide I frequency is very sensitive to the chemical environment, it can distinguish between K^+^ ions and water near the carbonyl group.^55, 59, 60^ To obtain site-specific information, two KcsA channels with Val76 and Gly77 residues ^13^C^18^O isotope labeled were synthesized. ^13^C^18^O isotope labeling shifts the amide I frequency by -66 cm^-1^ and, thus, resolves its vibrational signal from the majority of the unlabeled bulk signal.^52^ To obtain dynamical information at binding sites formed by Val76 and Gly77 residues, 2D IR experiments with 100 – 2000 fs waiting times between pump and probe pulses were performed. The change of the chemical environments of a vibrational chromophore during the waiting time typically leads to decorrelation of the frequencies in a process called “spectral diffusion”.^52^ If at short times 2D IR spectra are elongated along the diagonal, the spectrum is said to be inhomogeneously broadened and the distribution of frequencies typically corresponds to the distribution of chemical environments of a vibrational chromophore. Typically, 2D IR spectra become rounder as the waiting time increases, which is a measure of spectral diffusion. Spectral diffusion dynamics describes the time scales for frequency fluctuations which, in turn, reports on the time scales of structural dynamics.^61^ For the carbonyl groups in the selectivity filter, the time scale for frequency fluctuations is set by their interaction with K^+^ ions, water, and the surrounding protein.

Vibrational spectroscopy does not directly probe structure. Structural information can only be inferred with the help of theoretical models and simulations. We use molecular dynamics (MD) simulations and line shape theory to simulate 2D IR spectra for experimental waiting times. Simulations were performed for the two configurations [S1,W,S3,W] and [W,S2,W,S4] (Figure 1C), where “W” denotes water and “S1(2,3)” denotes a K^+^ ion occupying S1-S3 binding site. These configurations, as we recently showed, dominate in the selectivity filter of the KcsA channel in the closed conductive state.^62^ Here we found a very good agreement between experimental and simulated 2D IR spectra which enabled us to further use MD simulations to provide detailed information about the reorientation and hydrogen-bonding dynamics of water molecules in the selectivity filter as well as the fluctuations of K^+^ around the binding sites.

## RESULTS

Figure 2 shows experimental background subtracted (A-D) and simulated (E-F) 2D IR spectra of the Val76 isotope labeled KcsA channel for 100-2000 fs waiting times. The corresponding 2D IR spectra for the Gly77 labeled residue shown as well (J-Q). Figures S1 and S2 show the corresponding raw experimental spectra. The simulated spectra are for the mixture of the two configurations [S1,W,S3,W] and [W,S2,W,S4] at 3:2 ratio which, as reported in Ref.^62^, best fits both Val76 and Gly77 experimental spectra at a single waiting time of 500 fs. Here, for the first time, we illustrate that the agreement between our simulations and experiments is also good at other waiting times. The center lines are shown for each 2D IR lineshape as green lines. Figure 2I and R show the center line slopes from experiment and simulations as a function of the waiting time. Simulated and experimental center line slopes are nearly perfectly parallel suggesting that our simulations correctly reproduce spectral diffusion which in turn reports on the local dynamics near carbonyl groups. Importantly, the center line slopes exhibit almost no decay on picosecond timescales showing that there are no dynamical processes that cause the amide I frequencies to decorrelate on that timescale. In case of Gly77, experimental center line slopes are essentially independent of time. This means that the dynamics at that site is even slower than that of Val76. For a complete decorrelation of vibrational frequencies backbone carbonyls must experience all possible environments which is certainly not happening on the time scale of our experiments as confirmed by simulations. The good agreement between experimental and simulated 2D IR line shapes suggests that we can trust the accuracy of the MD simulations to obtain a more detailed picture of the dynamics inside the selectivity filter.

**Figure 2.**
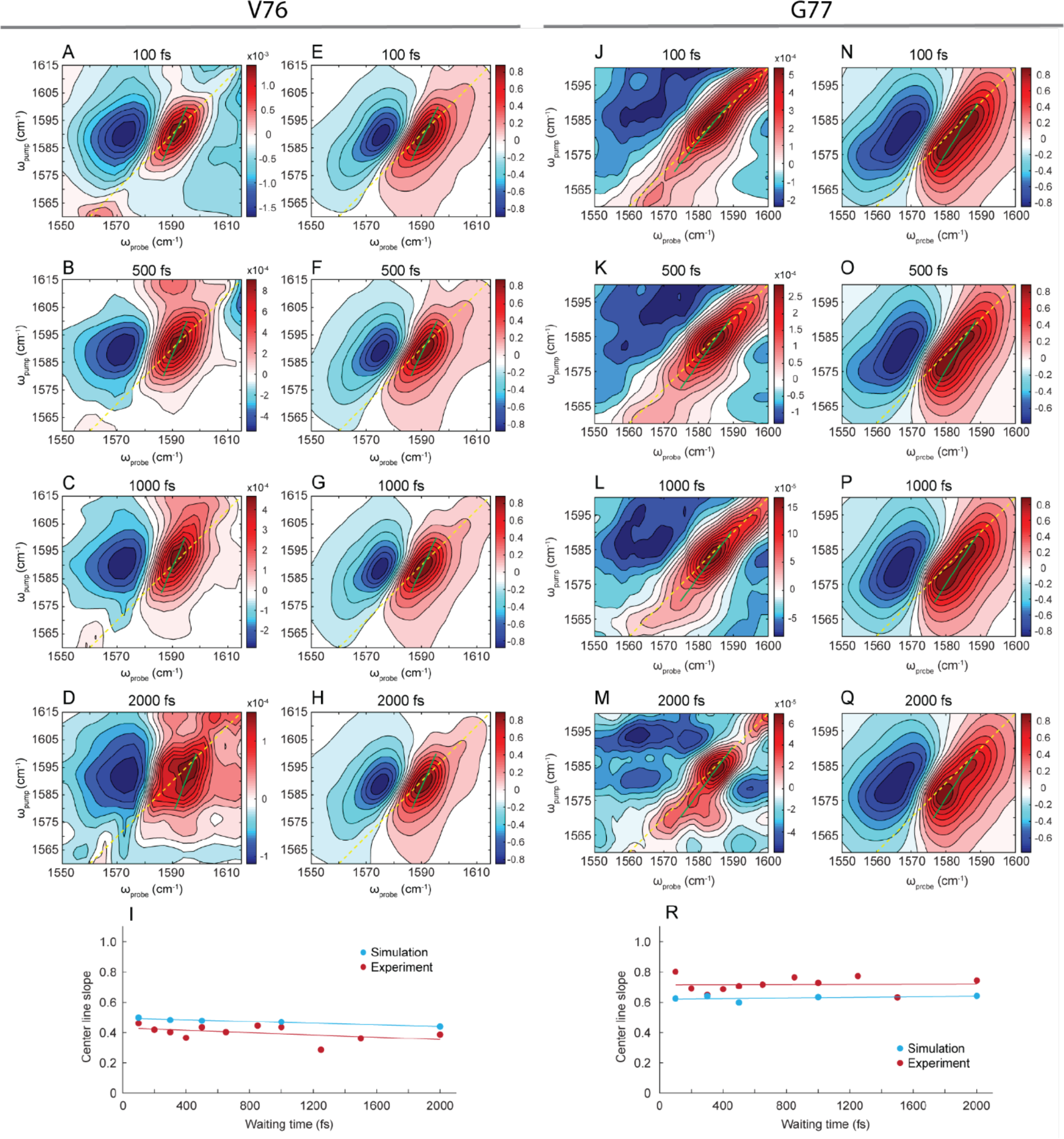
Experimental (A-D, J-M) and simulated (E-H, N-Q) 2D IR spectra of the KcsA channel with Val76 (A-H) and Gly77 (J-Q) isotope-labeled residues. Simulated spectra correspond to 60% of [S1,W,S3,W] and 40% of [W,S2,W,S4] ion configurations.^62^ Green lines show the center lines. Lower panels (I, R) show the center line slope of the experimental and simulated 2D line shapes as a function of time.

In what follows we analyze the dynamical properties of water molecules at the S1 site of the [W,S2,W,S4] configuration, water at the S2 site of the [S1,W,S3,W] configuration, and water at the S3 site of the [W,S2,W,S4] configuration. To indicate the location of the water being discussed, the corresponding “W” will be underlined “<u>W</u>”. For example, the water molecule at the S1 site of the [W,S2,W,S4] configuration will be denoted as [<u>W</u>,S2,W,S4]. We analyze frequency-frequency correlation functions (FFCFs, see Methods) of Val76 and Gly77 amide I vibrations, as well as the reorientational and hydrogen-bond dynamics of water. The reorientational dynamics of water will be described by the OH bond rotational time-correlation function defined as *C*_2_(*t*) = ⟨*P*_2_ (*û*(0) · *û*(*t*))⟩ where *û*(*t*) is the unit vector along the OH bond and *P*_2_ (*x*) = (3*x*^2^ − 1)/2 is the second-order Legendre polynomial. Here, the average is considered over both OH groups regardless of whether they form hydrogen bonds with the backbone carbonyl group or not. This rotational time-correlation function is a good approximation to the anisotropy decay measured in time-resolved IR pump-probe experiments.^3^ The timescales for molecular reorientation were obtained by fitting the rotational time-correlation function, after the initial decay, into a single exponential function.^63, 64^ Hydrogen-bonding dynamics can be quantified by the equilibrium hydrogen-bonding time-correlation function *C*_*hb*_(*t*) = ⟨*δn*(0)*δn*(*t*)⟩/⟨*δn*^2^⟩,^65^ where *n* is the number of hydrogen bonds that a water molecule makes with its neighbors, *δn*(*t*) = *n*(*t*) − ⟨*n*⟩, and is the average number of hydrogen bonds. The long-time decay of this function gives the characteristic time scales for the hydrogen-bond breaking and making dynamics. The geometric definition of the hydrogen bond introduced by Luzar and Chandler was used.^66, 67^

We separately analyze the contributions to the amide I 2D IR lineshape from protein, K^+^ ions, and water by calculating the respective FFCFs. CLS and FFCF are analytically related in certain limiting cases, but the FFCF provides a more detailed description of the vibrational dynamics. Moreover, FFCFs can be easily simulated by applying vibrational frequency maps to atomistic trajectories from MD simulations as described in Methods section. FFCFs calculated separately for protein, K^+^ ions, and water inside the selectivity filter for Val76 and Gly77 isotope labeled residues and each configuration are shown in Figure 3A. We note that water and K^+^ ion FFCFs were calculated by including the contribution of only water molecules and K^+^ ions, respectively, inside the selectivity filter. All other water molecules and K^+^ ions were excluded.

**Figure 3.**
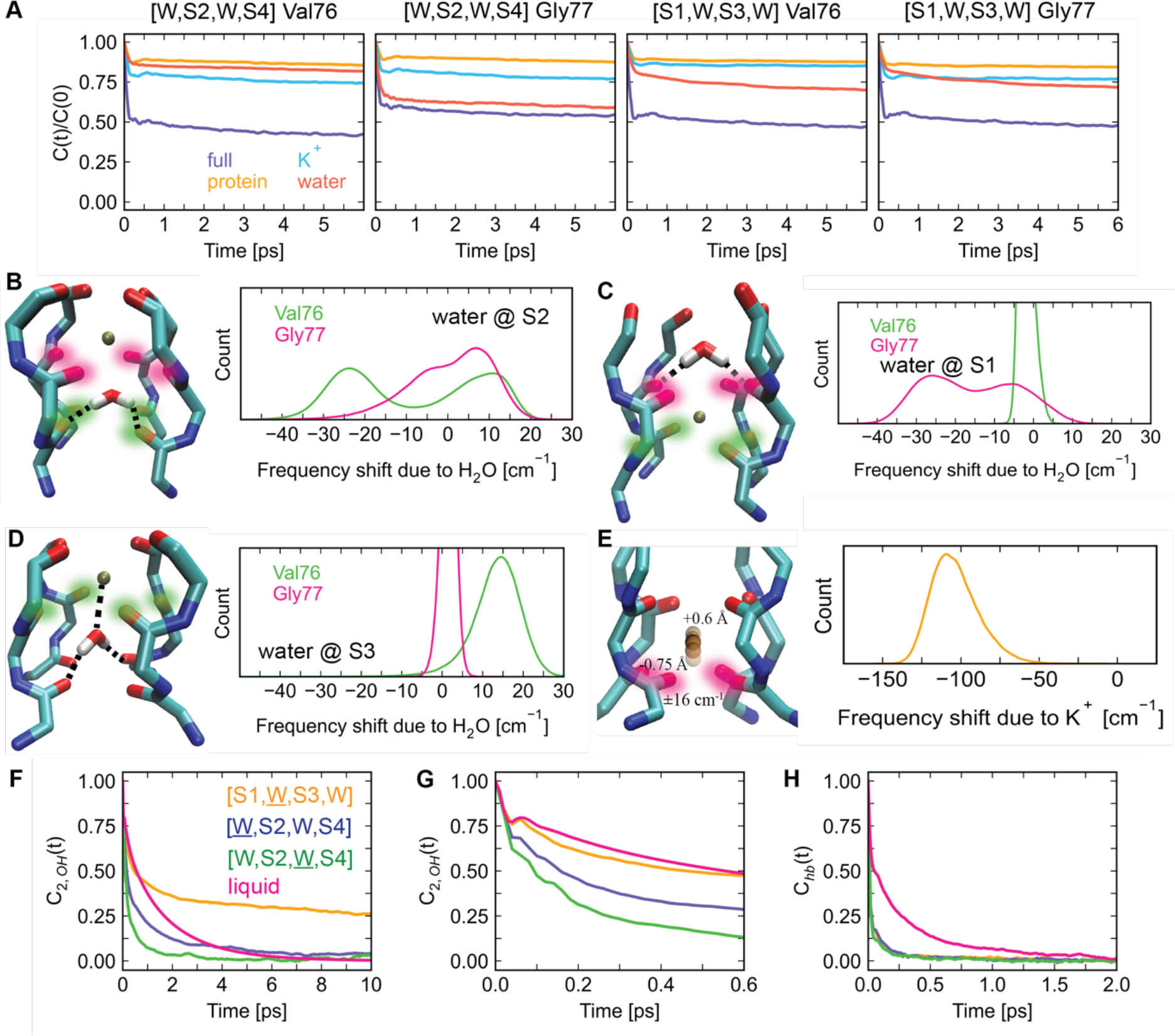
(A) Amide I frequency-frequency correlation functions (FFCFs) for Val76 and Gly77 residues and [W,S2,W,S4] [S1,W,S3,W] configurations. Each FFCF is an average over four FFCFs calculated for each amide I chromophore in a protein chain. (B-D) Distribution of amide I frequency shifts of Val76 (green) and Gly77 (pink) caused by water molecules and the structural fragments of the selectivity filter that are consistent with the frequency shifts and FFCFs. (E) Distribution of amide I frequency shifts caused by K^+^ ions and the schematic illustration of the magnitude of fluctuation of K^+^ ions that causes the observed frequency shifts. (F) Rotational time-correlation functions for water molecules at various sites in the selectivity filter compared to bulk liquid TIP4P water at 300 K (pink). (G) Zoom-in of the first 600 fs of rotational time-correlation functions showing the initial inertial decay. (H) Hydrogen-bond number fluctuation time-correlation functions.

All FFCFs exhibit a fast initial decay which causes motional narrowing. For protein and K^+^ ions’ FFCFs, the initial decay is followed by nearly static offsets which is indicative of Bloch dynamics.^68^ Water is the only component whose FFCFs continue to decay monotonically after the initial decay, which is especially pronounced for the [S1,W,S3,W] configuration. For the [W,S2,W,S4] configuration, water’s FFCFs decay slower, resembling the FFCFs of protein and ions. These results suggest that, unlike protein and K^+^ ions, water can be dynamic on a timescale of a few picoseconds. That being said, the dynamics of water is much slower than observed in the bulk.

Table 1 summarizes the amide I frequency shifts due to protein backbone, K^+^ ions, and water in the selectivity filter. K^+^ ions cause the largest frequency shift while water causes the smallest average shifts. Therefore, despite appreciable dynamics of water, the total amide I frequency shifts are dominated by K^+^ ions and the protein, with the corresponding FFCFs having Bloch dynamics. The static offset of the FFCF is related to structural inhomogeneity. These offsets are commensurate with the observed inhomogeneously broadened line shapes which are caused by K^+^ ions and the protein. We also note that the sum of these three contributions is within a few wavenumbers from the experimental peak frequencies (Table 1) which illustrates that i) other contributions (e.g., from lipid, water, and ions outside the selectivity filter) are negligible and ii) our approach to calculating vibrational frequencies is accurate.

**Table 1.** Mean amide I frequency shifts (cm^-1^) due to protein, water, and K^+^ ions in the selectivity filter and the total frequency shift from these contributions. The frequency shifts due to protein are the sum of the nearest-neighboring frequency shifts and the shifts due to electrostatic contributions from all atoms constituting local chemical environment of the protein (excluding nearest neighbors). Peak frequencies are determined by adding the frequency shifts to the ^13^C^18^O-labeled amide I frequency with no external field.^69^

| configuration<br>contribution | Val76 |  | Gly77 |  |
| --- | --- | --- | --- | --- |
|  | [W,S2,W,S4] | [S1,W,S3,W] | [W,S2,W,S4] | [S1,W,S3,W] |
| protein | 86.2 | 57.5 | 65.0 | 82.7 |
| water | -8.25 | 10.7 | 0.47 | -11.0 |
| $K^+$ ions | -109 | -90.2 | -90.9 | -103.6 |
| total frequency shift | -31.1 | -22.0 | -25.4 | -31.9 |
| peak frequency | 1587 | 1596 | 1592 | 1586 |
| experimental peak frequency | 1590 |  | 1585 |  |

We begin our analysis with the [S1,<u>W</u>,S3,W] water. The S2 site is bounded by Val76 and Gly77 residues. Therefore, this water can be expected to modulate the amide I frequencies of both Val76 and Gly77. The corresponding FFCFs (Figure 3A) clearly confirm that this is indeed the case. Figure 3B shows Val76 and Gly77 amide I frequency shifts due to the [S1,<u>W</u>,S3,W] water. The shifts are calculated for snapshots taken every 20 fs along a 2 ns long MD trajectory. We will use amide I frequency shift distributions to infer the structural arrangements of water in this and other cavities up to 2 ns (the timescale of our MD simulations). Figure 3B shows that the [S1,<u>W</u>,S3,W] water causes a -25 cm^-1^ frequency shift for Val76 carbonyls. This shift is distinct and is caused by hydrogen bonding between water and Val76 carbonyls. A positive frequency shift of +15 cm^-1^ is caused by Val76 carbonyls that are not hydrogen-bonded and face this water’s oxygen atom. Such a peak pattern can arise when the water molecule is hydrogen-bonded to the two nearest neighboring Val76 carbonyls. The geometry of the carbonyls, however, does not allow for two long-lived hydrogen bonds. This is supported by the hydrogen-bonding correlation function shown in Figure 3H. The hydrogen-bonding dynamics of the [S1,<u>W</u>,S3,W] water is an order of magnitude faster than in liquid water with the characteristic hydrogen-bond lifetime of just 60 fs compared to 0.8 ps calculated for the TIP4P water model.

Reorientational dynamics of the [S1,<u>W</u>,S3,W] water is illustrated by the corresponding correlation functions shown in Figure 3F-G. The initial fast decay is due to inertial rotational reorientation. One would expect a faster initial reorientational motion of the water molecules in the selectivity filter because of the fewer hydrogen bonds this water can form, leading to less hindered rotational and translation motions. Indeed, all rotational correlation functions for waters inside the selectivity filter initially decay faster compared to that of liquid water. The rotational correlation time for the [S1,<u>W</u>,S3,W] water is 3.3 ps which is slower than that of liquid TIP4P water (1.2 ps), in a good agreement with previously reported values for TIP4P water.^64^ Such short-lived hydrogen bonds and slow reorientation times paint the picture of this water molecule switching between being doubly hydrogen-bonded to next-neighboring Val76 carbonyls and singly hydrogen-bonded to one of the carbonyls all while staying approximately in the plane formed by the Val76 carbonyls. Remarkably, two clear peaks in the frequency shift distribution suggest that this configuration is stable on the timescales of our MD simulations. For example, this water does not trade its Val76 carbonyls as hydrogen-bond partners for Gly77 carbonyls. If this water molecule was dynamically exploring configurations within the cavity on picosecond timescales the frequency shift distributions of Gly77 and Val76 would be very close. Our atomistic interpretation of the configuration of this water is schematically shown in Figure 3B.

We now focus on [<u>W</u>,S2,W,S4] water. Because the Val76 residue is further down the protein chain (Figure 1B), water at the S1 site does not contribute significantly to Val76 amide I frequencies, which is corroborated by the very slow decay of the corresponding FFCFs ([W,S2,W,S4] Val76 plot in Figure 3A). In contrast, this water modulates the Gly77 amide I frequency more strongly than the protein and K^+^ ions. The [<u>W</u>,S2,W,S4] water has a rotational correlation time of 1.0 ps which is close to that of liquid water. The hydrogen-bond lifetime for this water is 60 fs, the same as for the [S1,<u>W</u>,S3,W] water. The amide I frequency shift distribution for Gly77 has the characteristic hydrogen-bonding peak at -25 cm^-1^. The absence of a positive frequency shifts indicates that there is no configuration in which the [<u>W</u>,S2,W,S4] water orients its oxygen towards Gly77 carbonyls. This observation, together with the faster relaxation of this water compared to the [S1,<u>W</u>,S3,W] water can be rationalized if this water molecule forms hydrogen bonds with non-adjacent Gly77 carbonyls. Similarly to the [S1,<u>W</u>,S3,W] water, the hydrogen-bonding geometry is imperfect, forcing this water molecule to rattle between the non-adjacent Gly77 carbonyls. This configuration is also found to be stable throughout the entire 2 ns molecular dynamics trajectory. Figure 3C shows the [<u>W</u>,S2,W,S4] water configuration that is consistent with the aforementioned results.

The [W,S2,<u>W</u>,S4] water does not form hydrogen bonds with Val76 carbonyls, which delineate the S3 site, on picosecond-to-nanosecond time scales. If hydrogen-bonded, this water would be expected to impact the amide I frequency of Val76 residues. However according to the corresponding FFCF (leftmost panel in Figure 3A), water’s contribution to the Val76 amide I frequency dynamics is slow and similar to that of the protein and K^+^ ions. Further evidence for this assignment comes from the corresponding distribution of amide I frequency shifts shown in Figure 3D. A single pronounced peak at a positive frequency shift means that this water molecule is oriented with its oxygen atom towards the plane formed by Val76 carbonyls during the whole 2 ns MD trajectory. The hydrogen bond lifetime for this water is the shortest and is only 40 fs. The rotational relaxation time of the [W,S2,<u>W</u>,S4] water is the fastest with a correlation time of only 0.40 ps. The average distance from this water’s oxygen atom and the K^+^ ion at the S2 site is 2.75 Å which is noticeably shorter than the distance between the water molecule at S1 and the K^+^ ion at S2 (3.43 Å). Interestingly, this value corresponds to the first peak of the K^+^-O radial distribution function (in bulk water) at 2.73±0.05 Å.^70^ Thus, this water molecule is oriented in the cavity such that its oxygen atom is coordinated by a K^+^ ion, and the water molecule resides at the most probable distance a water molecule can be found around a K^+^ ion in solution. The fastest reorientation and hydrogen-bond times across all studied configurations indicate that this water molecule might only be very weakly hydrogen-bonded to Thr75 carbonyls. Overall, this water molecule resembles “gas-phase” water (Figure 3D). Remarkably, this configuration is also stable. Despite being effectively free, within the confinement of the cavity, this water remains gas-phase-like throughout the entire molecular dynamics trajectory. This is likely because the attractive electrostatic forces between this water, K^+^ ion, and Thr75 carbonyls and the repulsive electrostatic forces due to Val76 carbonyls finely balance each other to furnish this water with the gas-phase-like behavior.

FFCFs indicate that K^+^ ions do not move appreciably, but they can still fluctuate within their binding sites. One can use amide I frequencies to understand the magnitude of K^+^ ion fluctuations. Figure 3F shows the distributions of amide I frequency shifts due to K^+^ ions. The standard deviations of the distributions are 23 cm^-1^ for the [S1,W,S3,W] Val76, 16 cm^-1^ for the [S1,W,S3,W] Gly77, 18 cm^-1^ for the [W,S2,W,S4] Val76, and 15 cm^-1^ for the [W,S2,W,S4] Gly77 configurations. We can estimate the magnitude of the fluctuation of K^+^ ions because electrostatic frequency maps relate frequency shifts to distances between K^+^ ions and C and N atoms of the peptide group (see Methods section). Consider, for example, the [S1,W,S3,W] configuration with the K^+^ ion placed at the S1 site exactly at the center of mass of the four Gly77 and four Tyr78 carbonyls that define the boundary of the ion binding site (Figure 3E). For a ±16 cm^-1^ frequency shift, the K^+^ ion needs to move ∼0.61 Å away from its position towards Tyr78 carbonyls or ∼0.75 Å closer to Gly77 carbonyls. These magnitudes of ion fluctuations estimated from frequency shifts are in agreement with previous MD simulations.^71^ To put this value into perspective, the distance between Gly77 and Tyr78 carbonyls that create the S2 site is ∼3.5 Å. Therefore, to cause the simulated frequency shifts ions can fluctuate about 35-45% the distance to either carbonyl from the geometric center of a binding site.

## DISCUSSION AND CONCLUSION

According to the soft knock mechanism of ion transport, K^+^ ions cannot advance through binding sites without concerted motion of the intervening water. Consequently, the dynamics of water play a role, since hydrogen bonds between the water and backbone have to be broken and reformed, should they be present. Thus, knowledge about the dynamics of water inside the selectivity filter under equilibrium and non-equilibrium (i.e., with transmembrane potential) conditions is an important part of understanding the ion transport. Slow spectral diffusion observed in our 2D IR experiments and supported by line shape simulations indicates that there are no dynamical processes at the S1-S3 sites inside the selectivity filter that occur on a 2 ps timescale for the KcsA channel in the closed conductive state; water remains either hydrogen bonded or free. Simulations put the exchange time at longer than 2 ns.

The selectivity filter of K^+^ channels is a very different environment for a water molecule than bulk water. Water is confined by the protein and K^+^ ions occupying the neighboring ion binding sites, but the binding site has too large of a volume to enable multiple stable hydrogen bonds to form. Thus, unlike in liquid, water in the selectivity filter cannot form four hydrogen bonds, nor can even two hydrogen bonds exist for as long as they do in liquid, due to the imperfect geometry of carbonyl groups that form the binding pocket. Out of many possible types of water in the selectivity filter schematically shown in Figure 4, in this work three types were identified through molecular dynamics simulations. The first type is water that is hydrogen-bonded to two adjacent carbonyls (Figure 4.1). Compared to liquid water, the reorientational dynamics of this water is slower while hydrogen bonds are shorter-lived because liquid water provides a perfect tetrahedral coordination environment. The second type is water that is hydrogen bonded to non-adjacent carbonyls of the same plane (Figure 4.2). Such water molecules rattle between the two carbonyls competing to form hydrogen bonds with water because the geometry of non-adjacent carbonyls does not allow for the simultaneous formation of two strong hydrogen bonds. Consequently, the rotational and hydrogen bonding dynamics of this water are faster than in the bulk. Finally, “gas-phase-like” water with fast, sub-picosecond reorientation times has been observed as well (Figure 4.8). Such water was found to reside at the first peak of the K^+^-O (in bulk water) radial distribution function. Remarkably, the interconversion between these types of water does not occur during the 2 ns trajectory of our molecular dynamics simulations.

**Figure 4.**
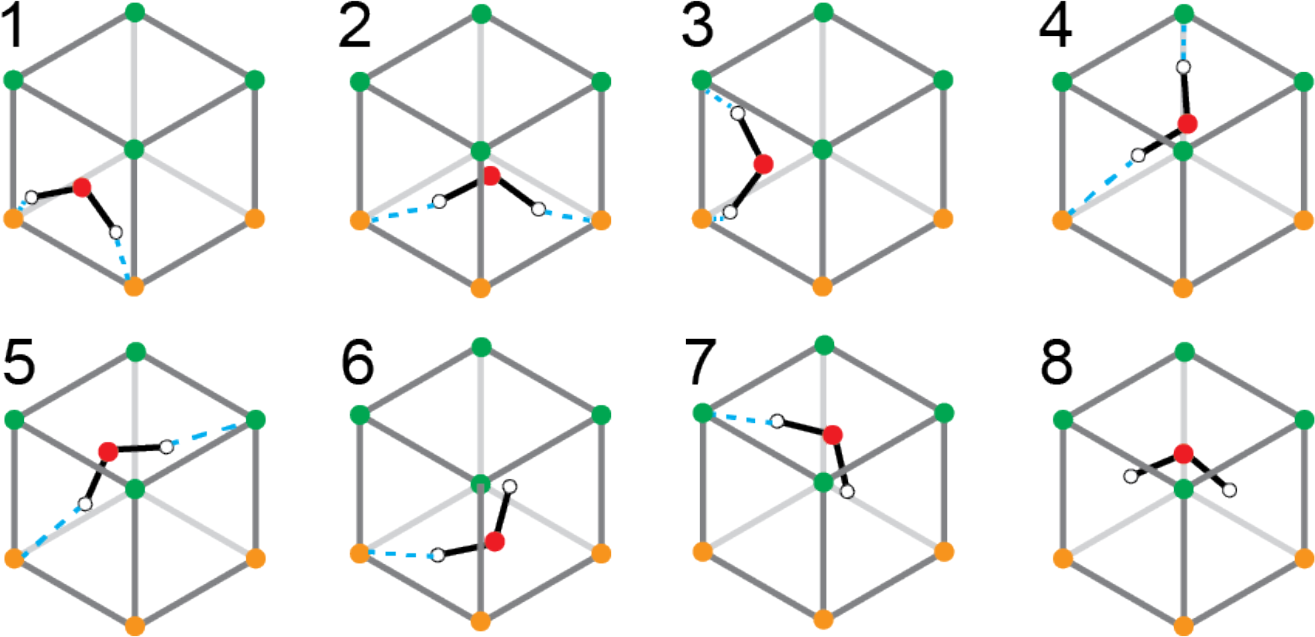
Schematic illustration of all possible ways a water molecule can be oriented inside a binding site cavity demarked by the two sets of four carbonyl groups colored in green and orange. Water’s oxygen atoms are shown in red, hydrogen atoms in white. Possible hydrogen bonds are indicated by blue dashed lines.

The water dynamics observed here are not typical of hydration water that operates as an interface between proteins and the bulk solvent. It is also not like structural water residing deep within a protein in well-defined locations, such as occurs when water acts as a proton shuttle. If anything, water in the selectivity filter of KcsA is more akin to water in non-biological samples like the pores of metal-organic frameworks (MOFs). MOFs are used as chemical filters by selecting the pore size and functional groups that line the pores. Gaseous water can diffuse into the pores and be stabilized by the functional groups but can also be remain non-bonded and freely spinning, similar to the different modes we observe for KcsA.^72, 73^ Perhaps the binding pockets of KcsA are set not only to interact optimally with KcsA, but also to provide an environment where water is only weakly hydrogen bonded or even unconstrained for long periods of time and thereby not the limiting factor for ion diffusion.

## MATERIALS AND METHODS

### Isotope labeling of KcsA

KcsA was ^13^C^18^O isotope labeled at the Val76 and Gly77 sites using protein semisynthesis as previously described in detail.^28, 62^

### 2D IR measurements

The details of the mid-infrared light generation and 2D IR spectrometer used to collect 2D IR spectra have been described elsewhere.^74^ Briefly, a 40 W Yb:KGW regenerative amplifier (Light Conversion Carbide) at a 100 kHz repetition rate is used to pump an optical parametric amplifier (Light Conversion Orpheus Mid-IR). The output signal and idler beams are combined into a non-colinear GaSe difference frequency generation setup (Light Conversion Lyra). The mid-IR light had a pulse energy of 3 μJ and was centered at a wavelength of 6.2 μm. This light is used in a 2D IR spectrometer with a pump-probe geometry. A ZnSe beamsplitter is used to create the pump and probe lines by splitting the light in a 75:25 ratio. The mid-IR The pump-pulse pair is generated using a germanium acousto-optic modulator (AOM). The pump and probe pulses are overlapped in time and space and are focused on the sample using 90° off-axis parabolic mirrors. The probe and resulting colinear signal are directed into a monochromator (Princeton Instruments). The dispersed light is then detected using a 64-pixel mercury cadmium telluride (MCT) array detector (Infrared Systems). A 75 or 150 groove/mm (g/mm) diffraction grating is used in the monochromator. The probe axis resolution is 4 cm^-1^ with the 75 g/mm grating and 2 cm^-1^ with the 150 g/mm grating. The pump and probe axes were calibrated as previously described.^75^ Spectra were collected with a ZZZZ polarization scheme and at waiting times of 100, 200, 300, 400, 500, 650, 850, 1000, 1250, 1500, and 2000 fs.

To remove background signal in the ^13^C^18^O isotope label region, difference spectra are generated from control and isotope labeled samples. Control spectra are scaled using the peak intensity of the amide-II mode to correct for intensity differences. The scaling factor determined for the 500 fs waiting time is used for all spectra. Figures S1 and S2 show the control and labeled spectra before subtraction.

### Molecular dynamics simulations

We used molecular dynamics (MD) trajectories from Kratochvil *et al*. ^28^ The crystallographic structure of the KcsA channel (Protein Data Bank entry 1K4C^38^) using residues Ser22 to His124 was embedded in a 71 A x 71 A bilayer consisting of 117 dipalmitoyl-phosphatidylcholine (DPPC) lipid molecules. The system was solvated with ∼12500 water molecules forming a bulk liquid solution around the membrane. K^+^ and Cl^-^ ions were added to the bulk solution to neutralize the charge of the system and establish a concentration of 500 mM solution of KCl. K^+^ ions and water were placed based on locations of ion binding sites. A polarizable force field based on classical Drude polarizable models was used for the KcsA and DPPC.^76, 77^ This force field accounts for atomic polarizability by including the flexible point charge particles from heavy atoms.^78, 79^ K^+^ and Cl^-^ parameters that represent improved ion-protein interactions were taken from Li et al.^80^ SWM4 model was used to describe water.^81^ For each ion configuration [S1,W,S3,W] and [W,S2,W,S4] the simulation system was first equilibrated at constant pressure of 1 atm and temperature 298.15 K (NPT). This was followed by a production simulation at constant volume and temperature 298.15 K (NVT). The temperature of the system was maintained by coupling to a Langevin thermostat, with temperatures of the Drude particles maintained at 1 K. Electrostatic interactions were treated using the Particle Mesh Ewald (PME) method. Smoothing functions were applied to both electrostatic and van der Waals forces, with the cut-off of 12 A and switching function turn on at 10 A. A 5 kcal/mol harmonic restrained has been applied to the protein Ca atoms, excluding those at the selectivity filter. The selectivity filter residues (74 to 79) were constrained using a spring constant of 2000 kJ/mol/nm^2^. The coordinates of the K^+^ ions in each binding site of the selectivity filter were constrained with flat bottom potentials. Water molecules were not constrained. Approximately 100 configurations were generated for each simulation system at an interval of 20 fs between adjacent frames. NAMD2.10 package was employed in simulations using Drude polarizable force field.^82^ Each configuration is then employed as the starting structure for the spectroscopy calculations. Systems were first minimized using the steepest descent algorithm. Then a 20-ps long trajectory was generated by performing MD simulations at NVT ensemble with a 2 fs integration time step. GROMOS96 53a6 force field including SPC water model was used.^83, 84^ This force field was used to generate trajectories for 2D IR simulations because it was used to develop spectroscopic maps. SPC water model was used for water.^85^ This water model provides a good description of dynamical properties of water near proteins.^8^ The coordinates of all atoms saved every of 20 fs and used in spectroscopy calculations as described below. GROMACS 5.0.4 package was used in these simulations.^86^ The same trajectories were also for calculating rotational and hydrogen-bonding correlation functions. The total length of these trajectories is 2 ns.

### Calculation of vibrational frequencies and 2D IR spectra

2D IR spectra were calculated using mixed quantum-classical approach based on frequency and coupling maps.^55, 69, 87, 88^ The amide I frequencies were calculated using electrostatic frequency maps developed by Wang et al.^69^ and nearest-neighbor (NN) frequency maps.^89^ The former relate the frequency and *i*th amide I chromophore ω_*i*_ (in cm^-1^) to local electric fields: 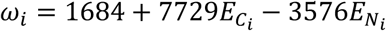, where 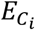 and 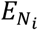 are the electric fields (in atomic units) in the C=O bond direction on the amide C and N atoms. These fields are due to nearby water, ions, peptide atoms that are more than 3 covalent bonds away from the peptide group, and all other atoms (e.g., lipids) within a cutoff of 20 Å. Electric fields are calculated from the point charges from GROMOS96 53a6 and SPC force fields. NN frequency maps account for the frequency shifts due to covalent through-bond effects from the nearest-neighbor amide groups. These frequency shifts depend on Ramachandran angles as explained in Ref. ^89^. The remaining contribution to amide I frequency is the shift due to ^13^C^18^O isotope label which was set to -66 cm^-1^. The vibrational anharmonicity is taken to be 14 cm^-1^ as determined experimentally in Ref.^90^. The lifetime of the first excited state of an isolated amide I chromophore was assumed to be independent of the residue and was set to 600 fs.^91^ In 2D IR calculations vibrational couplings between nearest-neighbor amide I chromophores were treated using nearest-neighbor coupling maps.^89, 92^ Longer-range couplings were treated using the transition dipole coupling scheme.^69, 93^ Amide I transition dipoles were calculated using the model of Torii and Tasumi.^92^ Frequency-frequency correlation functions were calculated for each amide I chromophore treating it is isolated, so the all couplings were set to zero. Frequencies, couplings, and transition dipole moments were calculated using MultiSpec program (https://github.com/kananenka-group/MultiSpec). 2D IR spectra in ZZZZ polarization are calculated using NISE program.^88^ The spectra are simulated with for same waiting times 100 – 3000 fs as in the experiment.

### Center line slope calculations

The center line slopes (CLSs) were determined as follows. For a given time *t* at each spectral cut along the ω _1_ frequency, the ω_3,*max*_ frequency is found where the spectrum peaks. CLS is then the slope of the resulted function. However, in practice, instead of calculating the derivative that is sensitive to the noise, it is useful to fit several (ω _1_, ω_3,*max*_) points in the spectral region of interest with a linear function and obtain the CLS value from the fit. The CLS has the maximum value of one corresponding to fully correlated the excitation and probing frequencies. As the slope with the increase of the waiting time T decays to zero, the correlation is completely lost.

### Frequency-frequency correlation function

Frequency-frequency correlation function (FFCF) is defined by *C*(*t*) = ⟨*δω*(0)*δω*(*t*)⟩, where *δω*(*t*) = *ω*(*t*) − ⟨*ω*⟩ is the instantaneous difference of the transition frequency from its average value.^52, 94-96^ FFCFs were calculated for Val76 and Gly77 amide I vibrations by treating each vibrational chromophore uncoupled from others, which is the common way of calculating FFCFs. Then the four FFCFs, one for each amide I chromophore in a protein chain, were averaged.

## Acknowledgements

Research reported in this publication was supported by the National Science foundation grant OIA-2229651 (A.A.K.) and National Institute Of General Medical Sciences of the National Institutes of Health under Awards R35GM150963 (A.A.K.) and R01GM135936 (M.T.Z. and F.I.V.). The content is solely the responsibility of the authors and does not necessarily represent the official views of the National Institutes of Health. A. A. K. also acknowledges the support from the start-up funds provided by the College of Arts and Sciences and the Department of Physics and Astronomy of the University of Delaware. Calculations were performed with high-performance computing resources provided by the University of Delaware.

## Supplementary Information

Experimental control and labeled 2D IR spectra before subtraction.

## Author contributions

M.T.Z. conceived the idea. All authors designed the research. M.J.R. performed 2D IR experiments. L.G. and F.I.V. synthesized the isotope-labeled KcsA channel. A.A.K. performed spectroscopy simulations and wrote the paper. A.A.K., M.J.R. and M.T.Z., analyzed the results.

## For Table of Contents Only

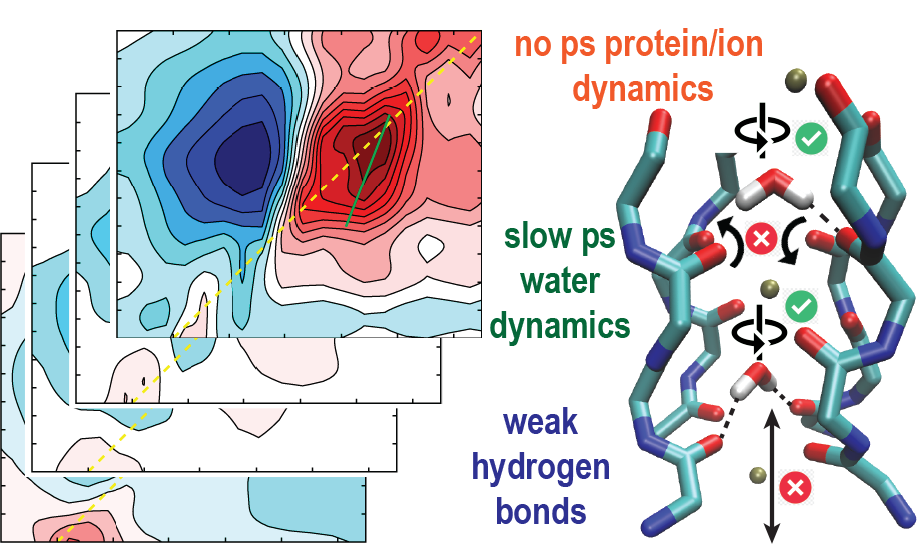

