## Supplementary Information for "Water inside the selectivity filter of a K^+^ ion channel: structural heterogeneity, picosecond dynamics, and hydrogen-bonding"

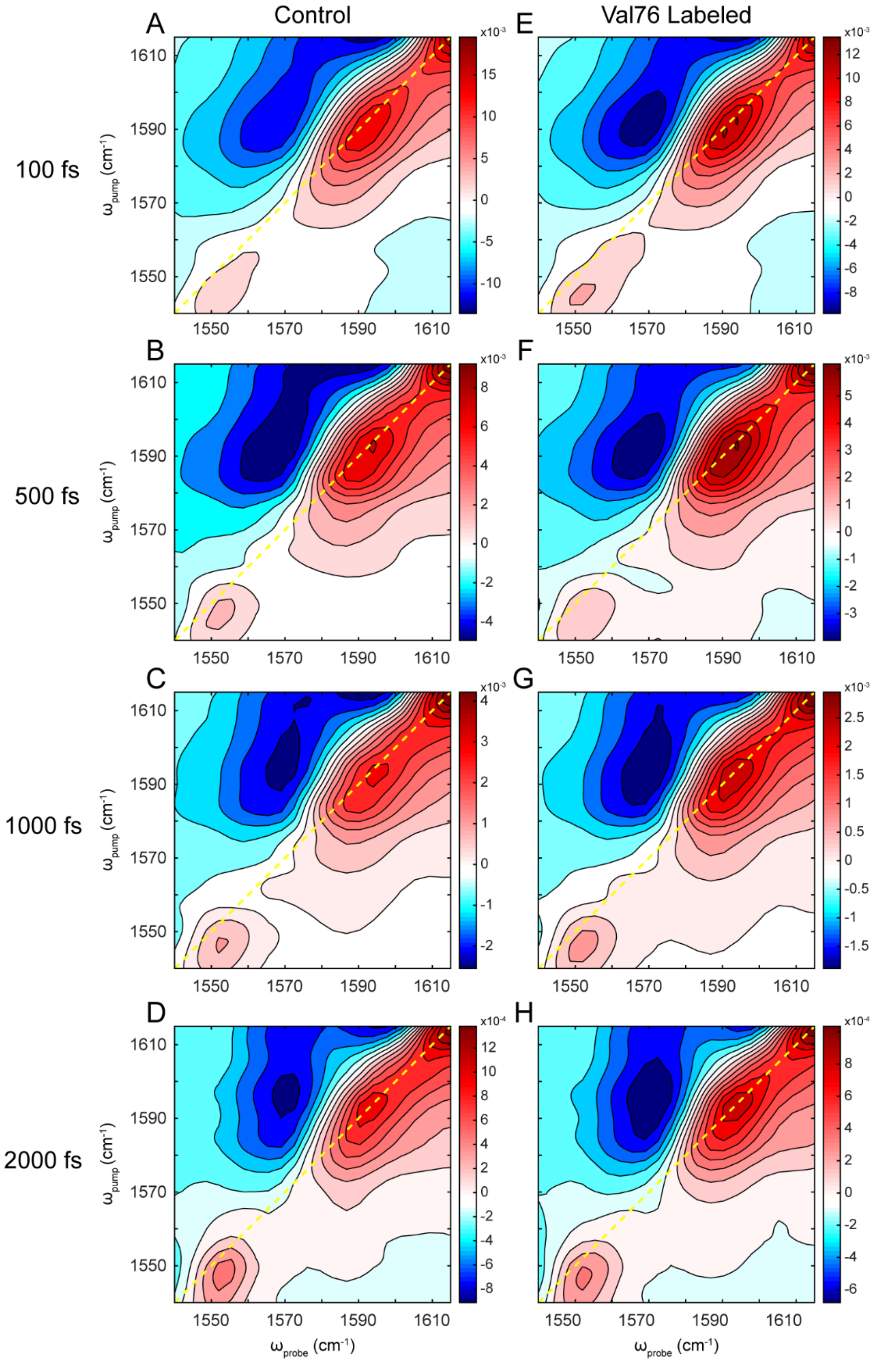


**Figure S1.** Experimental 2D IR spectra of the $^{13}C^{18}O$ isotope label region of control and Val76-labeled KcsA at various waiting times before normalization and subtraction. (A-D) are spectra of control (unlabeled) KcsA sample at 100, 500, 1000, and 2000 fs, respectively. (E-H) are spectra of Val76-labeled KcsA sample at 100, 500, 1000, and 2000 fs, respectively.

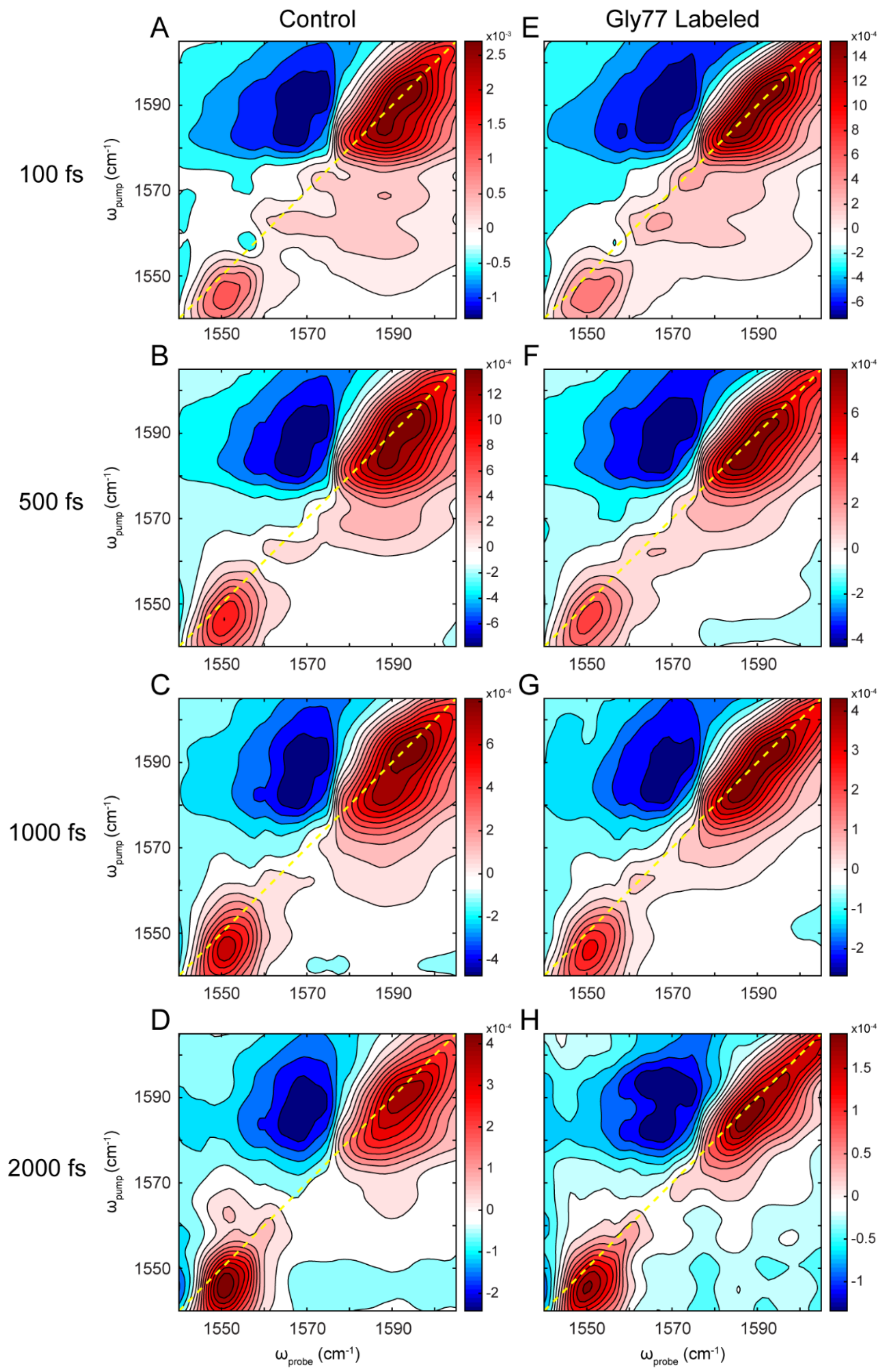


**Figure S2.** Experimental 2D IR spectra of the $^{13}C^{18}O$ isotope label region of control and Gly77-labeled KcsA at various waiting times before normalization and subtraction. (A-D) are spectra of control (unlabeled) KcsA sample at 100, 500, 1000, and 2000 fs, respectively. (E-H) are spectra of Gly77-labeled KcsA sample at 100, 500, 1000, and 2000 fs, respectively.
